# Changes in reward-induced neural activity upon Cafeteria Diet consumption

**DOI:** 10.1101/2023.10.20.563269

**Authors:** R. Heijkoop, J.F. Lalanza, M. Solanas, A. Álvarez-Monell, A. Subias-Gusils, R.M. Escorihuela, E.M.S. Snoeren

## Abstract

Excessive consumption of highly palatable foods rich in sugar and fat, often referred to as “junk” or “fast” foods, plays a central role in the development of obesity. The highly palatable characteristics of these foods activate hedonic and motivational mechanisms to promote food-seeking behavior and overeating, which is largely regulated by the brain reward system. Excessive junk food consumption can alter the functioning of this reward system, but exact mechanisms of these changes are still largely unknown. This study investigated whether long-term junk food consumption, in the form of Cafeteria (CAF) diet, can alter the reward system in adult, female Long-Evans rats, and whether different regimes of CAF diet influence the extent of these changes. To this end, rats were exposed to a 6-week diet with either standard chow, or *ad libitum* daily access to CAF diet, 30% restricted but daily access to CAF diet, or one-day-a-week (intermittent) *ad libitum* access to CAF diet, after which c-Fos expression in the Nucleus Accumbens (NAc), Prefrontal Cortex (PFC), and Ventral Tegmental Area (VTA) following consumption of a CAF reward of choice was examined. We found that all CAF diet regimes decreased c-Fos expression in the NAc-shell when presented with a CAF reward, while no changes in c-Fos expression upon the different diet regimes were found in the PFC, and possibly the VTA. Our data suggests that long-term junk food exposure can affect the brain reward system, resulting in an attenuated activity of the NAc-shell.

## 1. Introduction

As the prevalence of obesity continues to rise worldwide, its importance as a significant health concern grows. The condition can for example contribute to the development of cardiovascular disease, type 2 diabetes, hypertension and some cancers [1]. One of the key factors contributing to overweight and obesity is a lack of control over food intake [2], and large portion sizes and snacking patterns have been found to be risk factors in the development of obesity [3]. However, also excessive consumption of highly palatable foods rich in sugar and fat, often referred to as “junk” or “fast” foods, plays a central role in the development of obesity in humans [4]. The highly palatable characteristics of these foods activate hedonic and motivational mechanisms to promote food-seeking behavior and overeating [5], and it has been shown (also in rodents) that modern energy-rich diets are consumed not only because of their nutrient value but also because they are hedonically pleasant [6, 7].

The brain reward system has a central role in regulating motivation and reward-related responses. In normal circumstances, naturally rewarding behaviors like feeding, but also evaluation of palatability of the food, are regulated in part by dopamine transmission from the ventral tegmental area (VTA) to the nucleus accumbens (NAc) [8, 9]. Acute exposure to palatable foods increases dopamine synthesis and release [10, 11]. This increase in dopamine release, especially in the NAc, appears to play a central role in differentiating and encoding rewarding stimuli [12], resulting in an increase in the drive to engage in food seeking [13].

Excessive junk food consumption, however, can alter the functioning of this reward system [14-17]. The exact mechanisms of these changes are still largely unknown, but some studies have shown that rats that are obesity-prone show higher levels of motivation to work for food rewards than obesity-resistant or control animals [18-21]. Others, however, found that rats on a high-fat diet show reduced interest in cues for reward and show attenuated operant responses for sucrose [22, 23] or cocaine rewards [24].

Despite the large body of evidence indicating a relationship between junk food consumption and the reward system, it remains unclear what factors contribute to changes in the reward system and subsequent overeating. Besides the palatability of food, also the element of choice plays an important role in overeating. It has been shown that the body weight of rats on a high-fat, high-sugar diet of free choice increases more than that of rats on a forced high-fat, high-sugar diet, probably caused by the consumption of more meals per day without compensating meal size [25]. More variety in a meal and the greater dietary variation is an important element as well. The presentation of a wide range of foods evokes over-eating, known as the “buffet effect” [26]. The experimental *Cafeteria (CAF) diet* consists of several unhealthy, often ultraprocessed, but tasty products that humans consume too (e.g. bacon, muffins and cookies), and has because of that multiple advantages over purified, homogeneous high-fat/high-sugar diets [27-30]. It combines the orosensory properties (such as smell and texture) and palatability of the foodstuffs that promote overconsumption, but it also contains the element of free choice when the different products are provided simultaneously [28]. This way CAF diet mimics a certain pattern of problematic human consumption, which makes it very suitable as experimental diet model to study the effects of junk food consumption on the reward system.

Interestingly, when access to CAF diet is limited to a few days per week (often referred to as intermittent access), increases in amount of CAF consumption have been found [31, 32]. However, rats on an intermittent access scheme show intermediate to no responses to sucrose consumption compared to rats on a full-time CAF diet [31]. When rats had access to a restricted amount of CAF diet (30% of calorie restriction) besides standard chow, they showed decreased levels of sucrose preference similar to rats with *ad libitum* access to CAF diet compared with control rats, but had an intermediate phenotype on total sucrose intake in the two-bottle preference test and on the number of trials initiated in the brief access licking test, and no difference in hedonic responses to sucrose in the taste reactivity test [33, 34]. Altogether, this suggests that different exposures to CAF diet could have different effects on the reward system.

Therefore, this study investigates whether long-term CAF diet consumption, in which rats can choose the food products they consume, can alter the reward system in female rats, and whether different regimes of CAF diet influence the extent of these changes. To this end, rats received a 6-week diet with either standard chow, or *ad libitum* daily access to CAF diet, 30% restricted but daily access to CAF diet, or one-day-a-week (intermittent) *ad libitum* access to CAF diet. Subsequently, all rats were exposed to either a standard chow or CAF reward (again with a choice of food products) in order to investigate c-Fos expression as a marker for neural activity in the NAc, PFC and VTA. Changes in calorie intake and biometric variables were also compared. We hypothesized that chronic, daily *ad libitum* access to CAF diet would reduce the neural activation upon a food reward in the NAc, VTA and PFC, and that restricted or intermittent access to the CAF diet would result in intermediate effects compared to control diet.

## 2. Methods

All animals were cared for according to an institutionally approved experimental animal protocol, following Spanish legislation. The experimental protocol was approved by the Generalitat de Catalunya (approval number DAAM 9978) and was carried out in accordance with the EU Directive 2010/63/EU for animal experiments.

### 2.1 Animals and design

A total of 40 female Long-Evans rats (average body weight circa 230g upon arrival at an age of 13 weeks) were obtained from Janvier (Le Genest-Saint-Isle, France). The rats were pair-housed in standard polycarbonate cages at a temperature of 21±1°C and a humidity of 50±10% under a light/dark period of 12 h (lights on at 08:00) during the whole experiment.

After one week of habituation to the animal facility, the rats were weighed and randomly assigned to five treatment groups (n=8), which at baseline did not differ in average body weight. Each group underwent a different 6-week dietary regimen (Table 1). At the beginning of the seventh week rats were exposed to a food reward before perfusion for c-Fos expression analysis in the brain.

**Table 1.** Experimental groups exposed to standard chow or cafeteria (CAF) diet and reward.

| Group | 6 week diet | Reward |
| --- | --- | --- |
| CTR-Chow | *ad libitum* access to standard chow and water | Chow |
| CTR-CAF | *ad libitum* access to standard chow and water | CAF |
| CAFchr | *ad libitum* access to standard chow and water and daily *ad libitum* access to CAF diet | CAF |
| CAFint | *ad libitum* access to standard chow and water + 1 day per week *ad libitum* access to CAF diet | CAF |
| CAFres | *ad libitum* access to standard chow and water + daily access to 30% CAF diet (relative to CAFchr) | CAF |

### 2.2 Diet procedure

All groups of rats had *ad libitum* access to standard chow (Teklad Global 18% Protein Rodent Diet 2018” (Harlan, Barcelona, Spain)) and tap water. In addition, the rats in the CAFchr and CAFint diet groups received a CAF mealconsisting of a combination of the following products: pork jowl, biscuit with pâté, biscuit with cheese, muffins, carrots, and “flam” (jellied sugared milk) (details of the products can be found in Table S1). The rats on CAFres diet received a meal that consisted of a 30% calorie-restricted CAF diet (relative to the energy intake consumed by the CAFchr group) besides their standard chow. Rats in the CAFint group had *ad libitum* access to a CAF meal only once per week, while they received standard chow on the other daysAll meals were prepared fresh daily, and the amount eaten each day was registered. As the food products were given simultaneously, the rats were able to choose what food products to consume.

The mean daily food intake (g) during the 6-week diet exposure per cage, and the nutritional composition of the standard chow and CAF diet products are shown in tables 2, 3 and-S1. Some food product labels reported salt as NaCl. These values were recalculated to Na to enable uniform analyses of all food products.

**Table 2.**
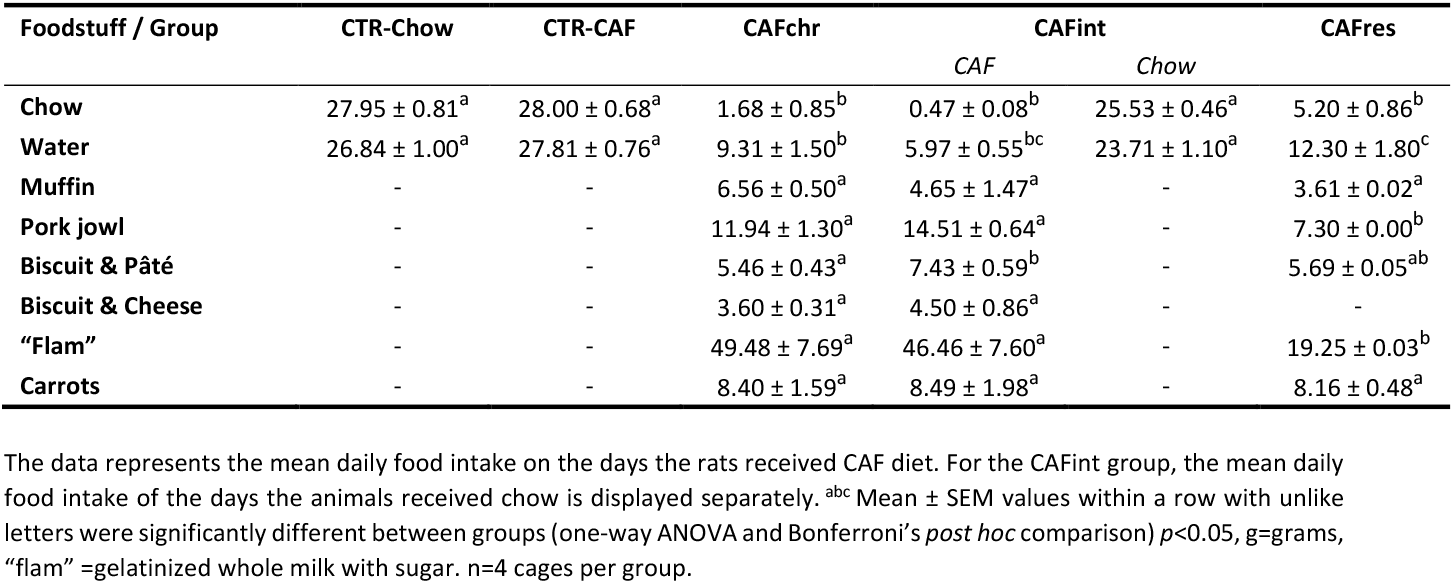
Mean daily food intake (g) during the 6-week diet exposure per cage.

**Table 3.** Mean daily intake of energy and nutrients during the 6-week diet exposure per cage.

| Nutritional Values / Group | CTR-Chow | CTR-CAF | CAFchr | CAFint |  | CAFres |
| --- | --- | --- | --- | --- | --- | --- |
|  |  |  |  | CAF | Chow |  |
| **Energy (kcal)** | 81.06 ± 2.35<sup>a</sup> | 81.19 ± 1.97<sup>a</sup> | 208.37 ± 1.99<sup>c</sup> | 224.28 ± 8.41<sup>c</sup> | 74.05 ± 1.34<sup>a</sup> | 125.69 ± 2.35<sup>b</sup> |
| **Proteins (g)** | 4.00 ± 0.12<sup>a</sup> | 4.00 ± 0.10<sup>a</sup> | 3.99 ± 0.17<sup>a</sup> | 4.05 ± 0.12<sup>a</sup> | 3.65 ± 0.07<sup>a</sup> | 2.74 ± 0.12<sup>b</sup> |
| **Carbohydrates (g)** | 13.42 ± 0.39<sup>a</sup> | 13.44 ± 0.33<sup>a</sup> | 18. ± 1.81<sup>b</sup> | 17.06 ± 0.71<sup>b</sup> | 12.26 ± 0.22<sup>a</sup> | 11.45 ± 0.38<sup>a</sup> |
| **Sugars (g)** | 0.00 ± 0.00<sup>a</sup> | 0.00 ± 0.00<sup>a</sup> | 14.30 ± 1.53<sup>c</sup> | 12.47 ± 0.54<sup>c</sup> | 0.00 ± 0.00<sup>a</sup> | 6.52 ± 0.04<sup>b</sup> |
| **Lipids (g)** | 1.12 ± 0.03<sup>a</sup> | 0.50 ± 0.01<sup>a</sup> | 12.70 ± 0.73<sup>c</sup> | 14.46 ± 0.69<sup>c</sup> | 1.02 ± 0.02<sup>a</sup> | 7.52 ± 0.04<sup>b</sup> |
| **Salt (Na) (g)<sup>1</sup>** | 0.0280 ± 0.0007<sup>a</sup> | 0.0280 ± 0.0006<sup>a</sup> | 0.0993 ± 0.0040<sup>a</sup> | 0.1086 ± 0.0054<sup>a</sup> | 0.0255 ± 0.02<sup>a</sup> | 0.0608 ± 0.0006<sup>b</sup> |
| **Water (g)** | 0.00 ± 0.00<sup>a</sup> | 0.00 ± 0.00<sup>a</sup> | 51.03 ± 5.87<sup>c</sup> | 173.10 ± 10.18<sup>c</sup> | 0.00 ± 0.00<sup>a</sup> | 24.98 ± 0.41<sup>b</sup> |

The data represents the mean daily food intake on the days the rats received CAF diet. For the CAFint group, the mean daily food intake of the days the animals received chow is displayed separately. <sup>abc</sup> Mean (± SEM) values within a row with unlike letters were significantly different between groups (one-way ANOVA and Bonferroni's *post hoc* comparison) $p < 0.05$ . kcal=kilocalories, g=grams. n=4 cages per group. <sup>1</sup> Values of salt content are presented as sodium (g).

A sample of daily food consumption was assessed once per week, except for the CAFint group which was assessed twice per week; once for CAF diet and standard chow, and once for standard chow only. Food consumption was calculated as the difference between the received amount of food product and the unconsumed amount of food product that was remaining 24 hours later.

### 2.3 Biometric variables

At the beginning of each week (Monday or Tuesday) during the 6-week diet exposure, body weight, abdominal perimeter, and length (nose-anus distance) were measured. From the combination of body weight and length we calculated the body mass index (BMI: body weight (g) / ano-nasal length (cm)^2^) and the Lee index (cube root of body weight (g) / length (cm)), which is an adaptation of the BMI for rodents [35].

### 2.4 Reward exposure

After six weeks of their respective diets, the rats were exposed to a food reward (CTR-Chow received chow, while the other groups received CAF with the freedom to choose different food products).. On the day of the reward exposure, the food was removed from the home cage at 07:00 am for short-term food deprivation. This was done to motivate rats to eat the reward. Approximately two hours before transcardiac perfusion, the rats were single-housed and exposed to 15% of the CAF diet or STD chow quantity (depending on the experimental group). The rats had access to the reward for 30 minutes (counted from the start of eating), before the remaining food was removed from their cages. Perfusion took place 90 minutes after removal of the food. No significant differences in latency to start eating were found between groups. Only one cage took more than 30 min to start eating and was therefore re-tested (and perfused) at the end of the day. Perfusions took place on three consecutive days and cages and diets were randomly assigned to each day and time of the day, balancing the experiment group.

For each rat, the amount of eaten reward products was collected the respective nutritional value of those products was calculated. The group results are presented in Supplemental Table S2 and S3, respectively.*2.5 Transcardiac perfusion*

The rats were deeply anesthetized with sodium pentobarbital (Doletal ®, 200 mg/kg i.p.) and then transcardially perfused with 0.1M phosphate-buffered saline (PBS; pH 7.4) followed by formaldehyde (3.7-4% buffered to pH=7 and stabilized with methanol <=3%, Casa Alvarez, Madrid, Spain). The brains were removed and post-fixed in individual vials with 4% formaldehyde for 48 hours. Subsequently, brains were transferred to 10% sucrose in 0.1M PBS solution, followed by a 20% and then a 30% sucrose in 0.1M PBS solution until they had sunken. A preservative (ProClin 150, Sigma-Aldrich, cat. 49377, 1:1000) was added to the final sucrose solution in which the brains were shipped to UiT The Arctic University of Norway for further analysis.

### 2.6 Brain sectioning and regions of interest

The brains were snap frozen in isopentane and kept at -80°C until sectioning. Brains were sectioned on a vibratome (Leica VT1200, Leica Biosystems, Nussloch, Germany) into 40 μm thick sections and stored in cryoprotectant solution (30% sucrose w/v, 30% ethylene glycol in 0.1 M PBS, pH 7.4) until further use. The brain regions of interest, of which the coordinates were determined using a rat brain atlas[36], were the nucleus accumbens shell and core (NAc-sh, NAc-c: AP between +2.28mm and +0.98mm), the anterior (aVTA) and posterior ventral (pVTA) tegmental area (-AP between -4.68mm and -6.12mm), and the prelimbic and infralimbic parts of the prefrontal cortex (PrL, IL: AP between +3.72mm and +2.52mm).

### 2.7 Immunohistochemistry

Sections were washed in Tris-buffered saline (TBS, 0.1M) before and after incubations. First, sections were blocked in a TBS solution with 0.5% BSA and 0.1% Triton-X-100 for 30 min at room temperature. Then, sections of each brain area were incubated in primary antibody for c-Fos (monoclonal rabbit anti-c-Fos, Cell Signaling, cat mAb-9F6, 1:4000) for 24h at room temperature, followed by 24h at 4°C. The samples were then incubated in secondary antibody solution (polyclonal biotinylated goat anti-rabbit antibody, Abcam, cat. Ab6720, 1:400) for 30min at room temperature, and subsequently with avidin-biotin-complex solution (Vectastain Elite ABC-HRP kit, Vector Laboratories, cat. PK-6100) in TBS for 30min at room temperature. Finally, the brain sections were incubated with 3,3’-diaminobenzidine (DAB substrate kit (HRP), Vector Laboratories, cat. SK-4100) for 5 min. After final washes, they were mounted on glass slides in 0.01M PB, dehydrated in ethanol, cleared with Xylene (Sigma-Aldrich, cat. 534056) and coverslipped using mounting medium (Entellan, Sigma-Aldrich, cat. 1.07961).

### 2.8 c-Fos immune-responsive cell quantification

After coverslipping, the slides were loaded into an Olympus VS120 slide scanner microscope. Image scans were obtained for each section using a 10x objective (NA 0.40), automatic focus settings in single plane, and fixed exposure of 1.13 ms. Using Olyvia online database software, high resolution cropped images (5x digital zoom) of the brain regions of interest were downloaded and opened in ImageJ, and bilateral counting boxes of always the same size and location were drawn to fit into the regions of interest. The following sizes of the boxes were used: NAc-shell and NAc-core: 0.81 mm x 0.51 mm; aVTA and pVTA: 0.6 mm x 0.5 mm; PrL: 1.0 mm x 0.4 mm; IL: 0.5 mm x 0.4 mm. Because the shape and size of the VTA vary substantially along the anteroposterior axis, placement templates for each AP location were created in ImageJ, in accordance with VTA locations in the Paxinos & Watson brain atlas. Subsequently, AP coordinates of each VTA section were determined using the brain atlas and the corresponding templates were used to ensure correct placement of the ROI boxes. Examples of the placement of ROI boxes can be found in Figure S1.

On these photomicrographs, all stained cells inside the zones were manually counted using Cell Counter plugin in ImageJ. Stained cells that overlapped with the counting frame border were included only if they overlapped with the dorsal or the lateral border. Cells that overlapped the ventral or medial border were not counted. This was performed for 2-3 sections per brain area per rat. Each section was counted twice when less than 10% difference in cell count was obtained, and 3 times when more than 10% difference was observed. Then the average of the two counts with less than 10% count difference was taken as data point. Per rat and brain region, the average of the 2-4 sections was used for further analysis. The number of counted cells was divided by the surface area of the zone to obtain the outcome measure of number of c-Fos-IR cells per mm^2^.

### 2.9 Statistical analysis

Since the rats were pair-housed, food intake and subsequent nutritional data was calculated per cage and converted in a mean daily intake of the 6 weeks diet. The other parameters were calculated for each individual animal and per week.

First, the CTR-Chow and CTR-CAF groups were compared, after which the different diet regime groups were compared only to CTR-CAF. For the food intake and nutritional data, an independent t-test was used to compare CTR-Chow and CTR-CAF, while the experimental groups were compared to CTR-CAF with a one-way ANOVA (taking the results of the Levene’s test for homogeneity of variances into account), followed by a Bonferroni post-hoc analysis.

A repeated measures ANOVA was employed for comparison of the biometric variables between the groups. When Mauchly’s test of sphericity was below 0.05, Greenhouse-Geisser was used for further analysis. In addition, a one-way ANOVA was performed to compare the differences between groups for each week, followed by a Bonferroni post-hoc analysis.

For the c-Fos data, t-test analysis was employed to compare CTR-Chow with CTR-CAF, and one-way ANOVA analyses were used for the different diet regime comparisons, followed by a Bonferroni post-hoc analysis. A p-value <0.05 was considered significant. Finally, a Pearson’s correlation was performed for the intake of the different nutrients during the reward phase and the number of c-Fos expressing cells.

## 3. Results

### 3.1 Food intake and nutritional data

An overview of the intake of diet components and correlating nutritional values per experimental group is shown in tables 2, 3 and S1. As expected, rats in the CTR-CAF and CTR-Chow groups did consume the same amount of water (*t*_(6)_=--0.775, p=0.967, table 2) and chow (*t*_(6)_=-0.043, p=468), and thus took a same amount of mean daily energy during the 6-week exposure to the diets per cage. In contrast, all three CAF groups (CAFchr, CAFint, CAFres) consumed significantly less standard chow and water during six weeks than the CTR-CAF control group (chow: F_(4,19)=_424.052, p<0.001; water: F_(4,19)_=59.229, p<0.001, table 2), except when the rats exposed to an intermittent access to CAF diet (CAFint) received the standard chow and they ate and drank the same amount as the control group. The intake of energy and nutrients differed significantly over the groups (energy: F_(4,19)_=287.643, p<0.001; protein: F_(4,19)_=20.970, p<0.001; sugar: F_(4,19)_=85.683, p<0.001; F_(4,19)_=196.426, p<0.001; carbohydrate: F_(4,19)_=12.587, p<0.001; salt: F_(4,19)_=37.733, p<0.001; and water: F_(4,19)_=187.794 p<0.001. Of note, a post hoc analysis showed that CAFres had consumed more sugars, lipids and salt than the CTR-CAF control group, but less than CAFchr and CAFint (detailed comparisons can be found in table 3). As expected, no differences were found between CTR-Chow and CTR-CAF group (energy: *t*_(6)_=-0.043, p=0.967, protein: *t*_(6)_=-0.039, p=0.970, carbohydrate: *t*_(6)_=-0.043, p=0.967, salt; *t*_(6)_=-0.043, p=0.967, and lipid: *t*_(6)_=-0.047, p=0.964, table 3).

### 3.2 Biometric data

We confirmed that no differences in body weight (F_(1,14)_=0.031, p=0.862, Fig. 1A), abdominal perimeter (F_(1,14)_=0.231, p=0.638, Fig. 1B), and Lee index (F_(1,14)_=0.455, p=0.511, Fig. 1C) were found between the two control diet groups before they were exposed to either a standard chow or CAF reward. In these control groups, there was also an expected increase in body weight (F_(2.794, 39.121)_=42.667, p<0.001), abdominal perimeter (F_(5,70)_=2.871, p=0.020), and Lee index (F_(2.786,39.010)_=51.458, p<0.001) over the course of the 6 weeks of diet exposure, but no interaction effect of diet*weeks was found (body weight: F_(2.794,39.121)_=0.438, p=0.713; abdominal perimeter: F_(5,70)_=2.224, p=0.061; Lee index: F_(2.786,39.010)_=1.431, p=0.249).

**Figure 1.**
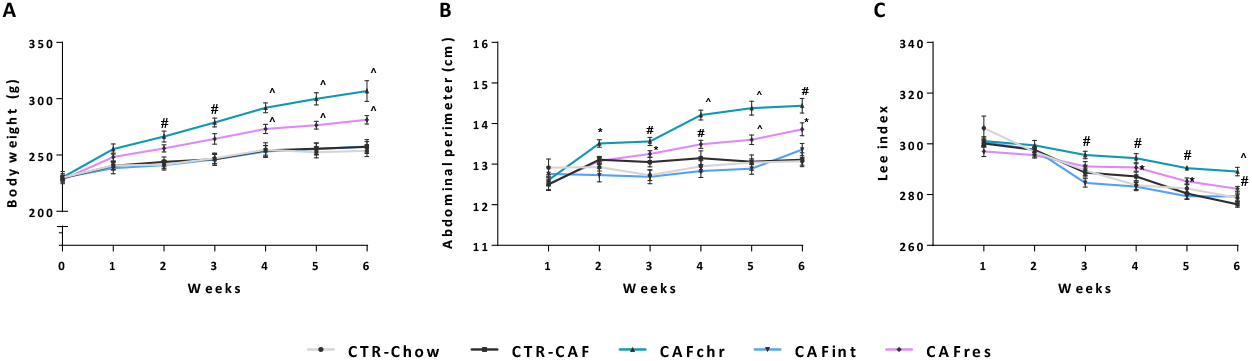
Biometric data of CTR-Chow (light grey line, n=8), CTR-CAF (dark grey line, n=8), CAFchr (green line, n=8), CAFint (blue line, n=8), and CAFres (purple line, n=8) female Long-Evans rats. (A) Body weight over the course of the 6 week diet exposure in grams. (B) Abdominal perimeter in centimeters. (C) Lee index (cube root of body weight (g) / ano-nasal length (cm)). a = p<0.05 significantly different from either CTR-CAF or CAFint (see results for details), b = p<0.05 significantly different from both CTR-CAF and CAFint, c= p<0.05 significantly different from all other groups.

When we compared the different CAF regimes to the control diet exposed to CAF-reward (CTR-CAF), we found an effect of diet and weeks on body weight (diet: F_(3,28)_=10.918, p<0.001; weeks: F_(2.984,83.562)_=225.375, p<0.001, Fig. 1A), abdominal perimeter (diet: F_(3,28)_=16.745, p<0.001; weeks: F_(3.584,100.352)_=63.631, p<0.001, Fig. 1B), and Lee index (diet: F_(3,28)_=10.462, p<0.001; weeks: F_(5,140)_=117.559, p<0.001, Fig. 1C). An interaction effect of diet*weeks was also found for body weight (F_(8.953,83.562)_=14.417, p<0.001), abdominal perimeter (F_(10.752,100.352)_=7.464, p<0.001), and Lee index (F_(15,140)_=4.353, p<0.001). Post-hoc analysis revealed that chronic *ad libitum* access to CAF (CAFchr) diet resulted in significantly higher body weights compared to control diet (CTR-CAF, p<0.01) or intermittent access (CAFint, p>0.001) to CAF diet, starting in the second week of exposure (Fig. 1A). From the second week of diet exposure, CAFchr also developed an increased abdominal perimeter compared to all other diet regimes (CTR-CAF: p<0.01, CAFint: p=0.010, CAFres: p<0.01). Restricted, but daily access to CAF diet (CARres) resulted in a significantly different abdominal perimeter than intermittent access to CAF diet (CAFint, p=0.032) from the third week as well (Fig. 1B). The Lee index were also significantly higher in CAFchr compared to CTR-CAF (p>0.001), CAFint (p>0.001), and CAFres (p=0.21) from the third week of exposure (Fig. 1C-D). Interestingly, intermittent access to CAF diet (CAFint) did not result in any different biometric outcomes compared to the control group (CTR-CAF).

### 3.3 c-Fos expression in the NAc, VTA and PFC

First, we tested whether acute exposure to a CAF reward induced c-Fos expression in the NAc, VTA and PFC in rats on control diet (standard chow) for 6 weeks. When we looked at the NAc shell, we found a significant increase in c-Fos expression upon an acute exposure to a CAF reward compared to a standard chow reward (*t*_(14)_=-3.080, p=0.008, Fig. 2A/C). We did not find these differences in the NAc core (*t*_(14)_=-0.828, p=0.422, Fig. 2B/D). No differences in number of c-Fos-IR (immunoreactive) cells were found between CAF and standard chow reward in the VTA (VTA anterior: *t*_(10)_=-1.429, p=0.092; or VTA posterior: *t*_(10)_=-0.276, p=0.394, Fig. 2E-H) or PFC (PrL: *t*_(12)_=-0.611, p=0.553; IL: *t*_(12)_=-0.213, p=0.425, Fig. 2I-L).

**Figure 2.**
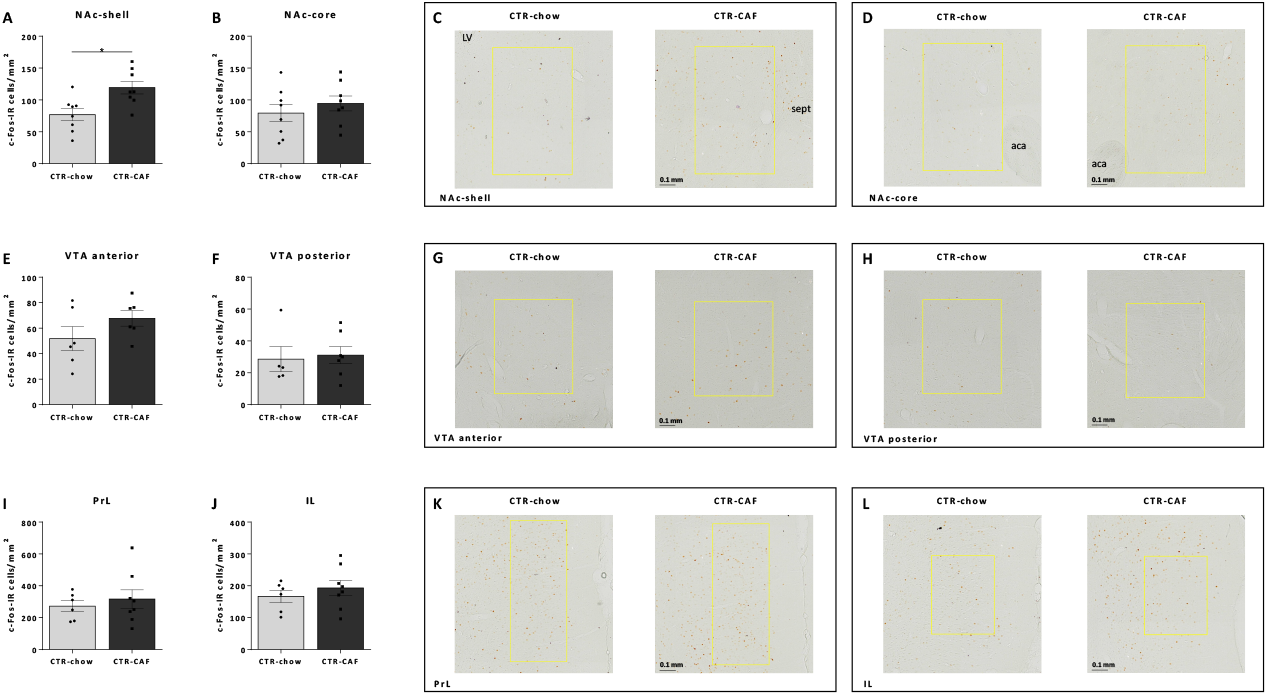
A) The total count of c-Fos-IR cells per mm^2^ in the nucleus accumbens (NAc)-shell and B) NAc-core, C) Examples of microphotographs with the counted c-Fos-IR cells of the different control groups in the NAc-shell and D) NAc-core, E) The total count of c-Fos-IR cells per mm^2^ in the anterior and (F) posterior ventral tegmental area (VTA), G)) Examples of microphotographs with the counted c-Fos-IR cells of the different control groups in the anterior and H) posterior VTA, I) The total count of c-Fos-IR cells per mm^2^ in the prelimbic prefrontal cortex (PrL) and (J) infralimbic cortex (IL), K) Examples of microphotographs with the counted c-Fos-IR cells of the different control groups in the PrL and L) IL. The yellow outlines represent the zone that was used for counting. The data are shown with individual data points, with the bars representing the mean ± standard error of the mean for CTR-chow (light grey bar, n=4-8) and CTR-CAF (dark grey bar, n=5-8) female Long-Evans rats. Aca= anterior commissure, LV = lateral ventrical, Sept = Septum. * p<0.05

Second, we studied the effects of different regimes of CAF diet on c-Fos expression upon exposure to a CAF reward. We found that long-term CAF diet (CAFchr) decreases the number of c-Fos-IR cells in the NAc-shell compared to a control diet of standard chow after exposure to a CAF reward (F_(3,29)_=11.839, p<0.001, Fig. 3A/B), but not the NAc-core (F_(3,29)_=2.182, p=0.114, Fig. 3C/D). A post-hoc analysis revealed that rats on a chronic *ad libitum* showed reduced levels of c-Fos expression in the NAc-shell (p<0.01), but not the NAc-core compared to CTR-CAF. With regard to the rats on a restricted CAF diet, a reduced c-Fos expression was found in the NAc-shell (p=0.004 compared to CTR-CAF). Rats on an intermittent CAF diet only showed reduced levels of c-Fos in the NAc-shell (p=0.013), compared to control rats that received CAF as reward.

**Figure 3.**
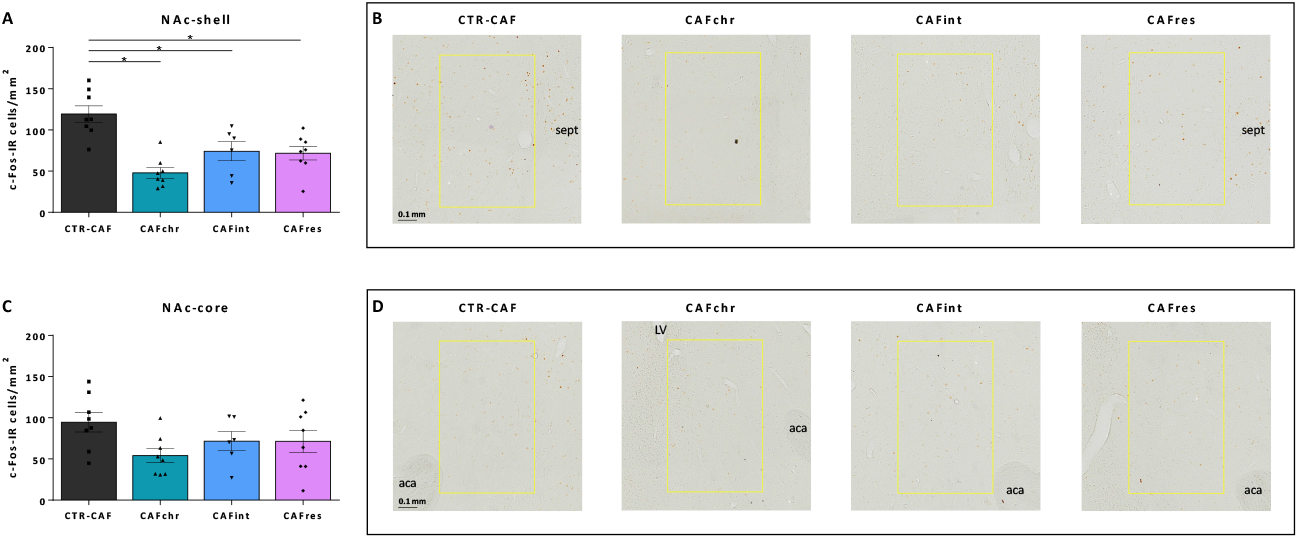
A) The total count of c-Fos-IR cells per mm^2^ in the nucleus accumbens (NAc)-shell, B) Examples of microphotographs with the counted c-Fos-IR cells of the different diet groups in the nucleus accumbens shell (NAc-shell,)C) The total count of c-Fos-IR cells per mm^2^ in the NAc-core, and D) the nucleus accumbens core (NAc-core). The yellow outlines represent the zone that was used for counting. The data are shown with individual data points, with the bars representing the mean ± standard error of the mean for CTR-CAF (dark grey bar, n=8), CAFchr (green bar, n=8), CAFint (blue bar, n=6), and CAFres (purple bar, n=8) female Long-Evans rats. Aca= anterior commissure, LV = lateral ventrical, Sept = Septum. * p<0.05

No differences in c-Fos expression between the diet regimes were found in the anterior (F_(3,23)_=2.179, p=0.122, Fig. 4A/B) or posterior (F_(3,23)_=0.339, p=0.797, Fig. 4C/D) VTA. At last, no differences in c-Fos expression between diet regimes were found in the PrL (F_(3,30)_=1.486, p=0.241, Fig. 5A/B) or IL (F_(3,30)_=1.201, p=0.328, Fig. 5C/D).

**Figure 4.**
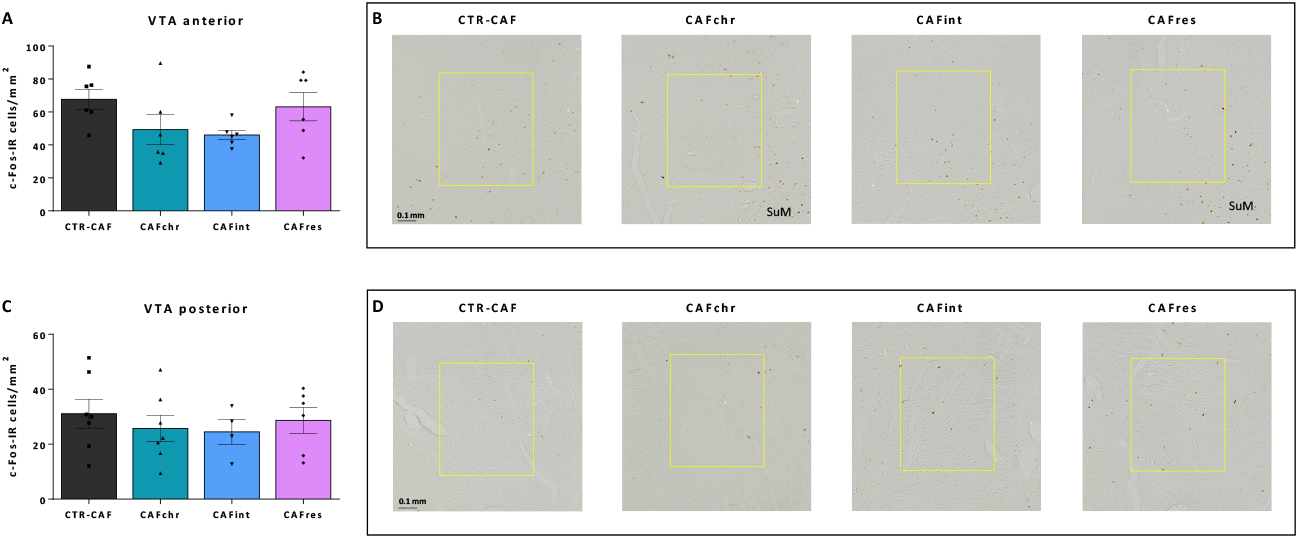
A) The total count of c-Fos-IR cells per mm^2^ of the anterior ventral tegmental area (VTA), B) Examples of microphotographs with the counted c-Fos-IR cells of the different diet groups in the anterior VTA, C) The total count of c-Fos-IR cells per mm^2^ in the posterior VTA, and D) Examples of microphotographs with the counted c-Fos-IR cells of the different diet groups in the posterior VTA. The yellow outlines represent the zone that was used for counting. The data are shown with individual data points, with the bars representing the mean ± standard error of the mean for CTR-CAF (dark grey bar, n=5-7), CAFchr (green bar, n=6-7), CAFint (blue bar, n=4-6), and CAFres (purple bar, n=4-6) female Long-Evans rats. SuM=supramammillary nucleus.

**Figure 5.**
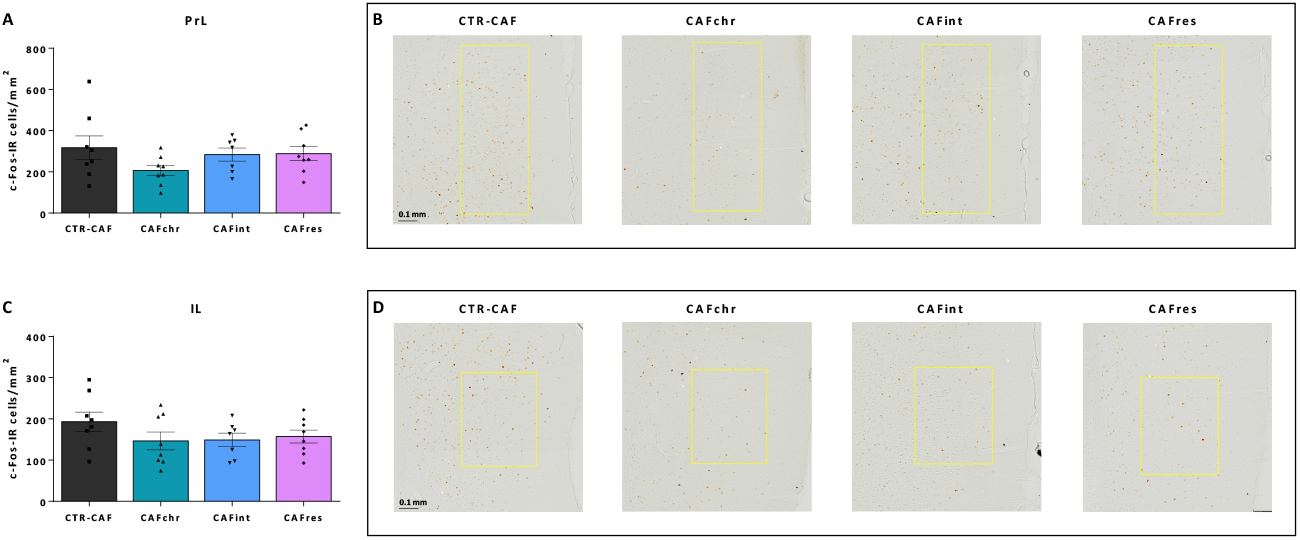
A) The total count of c-Fos-IR cells per mm^2^ in the prelimbic prefrontal cortex (PrL), B) Examples of microphotographs with the counted c-Fos-IR cells of the different diet groups in the PrL, C) The total count of c-Fos-IR cells per mm^2^ in the infralimbic prefrontal cortex (IL). and D) Examples of microphotographs with the counted c-Fos-IR cells of the different diet groups in the IL. The yellow outlines represent the zone that was used for counting. The data are shown with individual data points, with the bars representing the mean ± standard error of the mean for CTR-CAF (dark grey bar, n=8), CAFchr (green bar, n=8), CAFint (blue bar, n=7), and CAFres (purple bar, n=8) female Long-Evans rats.

Although the *choice* of food products was the most important element of the food reward stimulus, it is important to confirm that the eaten products do not have an influence on the c-Fos expression in the different brain regions. Analysis confirmed that neither energy (NAc-shell: ρ=0.036, p=0.828; NAc-core: ρ=-0.083, p=0.620; VTAa: ρ=0.103, p=0.587; VTAp: ρ=0.146, p=0.449; PrL: ρ=0.305, p=0.066; IL: ρ=0.136, p=0.421), protein (NAc-shell: ρ=0.007, p=0.967; NAc-core: ρ=-0.126, p=0.450; VTAa: ρ=0.063, p=0.739; VTAp: ρ=0.144, p=0.456; PrL: ρ=0.336, p=0.042; IL: ρ=0.136, p=0.421), sugar (NAc-shell: ρ=0.070, p=0.677; NAc-core: ρ=-0.053, p=0.752; VTAa: ρ=-0.023, p=0.905; VTAp: ρ=0.028, p=0.885; PrL: ρ=0.228, p=0.175; IL: ρ=0.024, p=0.887), lipids (NAc-shell: ρ=0.021, p=0.900; NAc-core: ρ=-0.066, p=0.696; VTAa: ρ=0.135, p=0.478; VTAp: ρ=0.150, p=0.438; PrL: ρ=0.286, p=0.086; IL: ρ=0.153, p=0.367), carbohydrates (NAc-shell: ρ=0.130, p=0.437; NAc-core: ρ=-0.064, p=0.704; VTAa: ρ=-0.104, p=0.584; VTAp: ρ=0.002, p=0.992; PrL: ρ=0.166, p=0.327; IL: ρ=-0.024, p=0.889), salt (NAc-shell: ρ=0.097, p=0.562; NAc-core: ρ=-0.057, p=0.736; VTAa: ρ=0.029, p=0.879; VTAp: ρ=0.136, p=0.482; PrL: ρ=0.100, p=0.557; IL: ρ=-0.052, p=0.759) nor water (NAc-shell: ρ=0.033, p=0.845; NAc-core: ρ=-0.086, p=0.608; VTAa: ρ=-0.211, p=0.264; VTAp: ρ=-0.067, p=0.729; PrL: ρ=0.162, p=0.337; IL: ρ=-0.072, p=0.671) had an effect on the number of c-Fos expressing cells in the NAc, VTA and PFC. On in the PrL, a small, though significant, positive correlation between the amount of eaten proteins and c-Fos expressing cells was found (ρ=0.336, p=0.042). This could have been cause by the rats on an intermittent CAF-diet who ate significantly more ‘flam’, which is high in proteins (Table S3). However, as this effect was not found back in the group differences, future research should reveal the role of proteins on neural activity in the PrL.

## 4. Discussion

In this study we investigated the effects of different long-term cafeteria diet regimes on the brain reward system in response to a food reward of choice. First of all, we found that an acute first-time exposure to a CAF reward caused an increase in c-Fos expression in the NAc-shell, but not in the VTA and PFC. In addition, we found that only chronic access to CAF diet (both restricted and *ad libitum*) resulted in higher biometric outcomes (body weight, abdominal perimeter, and Lee index) as other studies similarly reported [31, 37, 38]. However, chronic but also intermittent exposure to CAF diet caused a decrease in c-Fos expression in the NAc-shell upon a CAF reward compared to rats on a control diet. No differences in c-Fos expression levels were found in the PrL and IL of the PFC, nor in the aVTA and pVTA. The changes in c-Fos expression levels were not related to the nutritional intake of the reward of choice. These results suggest that regular junk food consumption, even if consumed only 1 day per week, can change the brain reward system, mainly by affecting the NAc-shell.

The increase in c-Fos expression in the NAc-shell upon an acute first-time CAF reward is in line with other studies that found increases in c-Fos expression in the NAc after drinking palatable glucose or sucrose [39, 40]. Although surprising at first sight, the lack of differences in c-Fos expression between a control and palatable food reward in the VTA and PFC has also been shown before [39]. However, in that study an increased neural activity has been found in other regions of the PFC (the ventrolateral and lateral parts) after drinking glucose [39]. Since dopamine synthesis and release increases following acute exposure to palatable foods [10, 41], and dopamine signaling plays a central role in differentiating and encoding rewarding stimuli [12], we expected that dopamine-rich brain regions like the VTA and dopamine-receiving areas like the PFC would have increased levels of neural activity, indicated by higher levels of c-Fos in those regions, in response to a CAF reward as well. As a tendency of increased levels of c-Fos-IR cells in the anterior VTA is seen, the lack of significant increases might in part have been caused by the anatomical sparseness of (dopamine) cells in the VTA and the limited number of brain sections that were available for this study.

All long-term CAF diet regimes did result in attenuated activity of the NAc-shell neurons in response to a CAF reward compared to rats on a control diet. That chronic *ad libitum* access to junk food can decrease neural activity in the NAc-shell is in line with another study that showed reduced levels of pERK after a chronic CAF diet when exposed to a sucrose solution [42]. In addition, many studies have shown that rats fed high in calorie food show lower dopamine turnover in the NAc at baseline or after a sucrose reward [22, 43-47]. This reduction in dopamine levels in the nucleus accumbens is suggested to be due to reduced stimulated dopamine release and vesicle size [45, 46]. In addition, it was found previously that rats on CAF or high-fat, high-sugar diets show reduced levels of ethanol or sucrose intake [48-50]. The diets seem to attenuate the rewarding properties of these substances, which could explain the lower levels of neural activation in the NAc-shell. It should be mentioned, though, that others have not been able to find changes in c-Fos expression in the NAc in rats on a high-fat diet, but this could be explained by differences in the duration of the palatable food exposure [39].

The interesting aspect of our study can mainly be found in the fact that also chronic restricted access to a CAF diet, and even 1 day a week exposure, already reduces c-Fos expression in the NAc-shell. One could argue that familiarity with the CAF reward may cause this decreased neural activation. The CTR-CAF rats have never been exposed to CAF before and could therefore experience the CAF reward as more rewarding. However, limited access to CAF has been shown to increase the amount of CAF consumption compared to chronic access [31, 32]. In addition, when exposed to a sucrose solution, rats exhibit more licks per cluster than rats on a chronic CAF diet [31]. If novelty plays a role in our finding, it would then be expected that also the CAFint rats would show increased, instead of decreased c-Fos expression in the NAc-shell. Either way, more research is needed to unravel the reasons behind these changes in NAc-shell neural activation.

As mentioned above, our findings surprisingly did not show an effect of exposure to an acute CAF reward on c-Fos-IR cell levels in the VTA in control rats. Due to unforeseen circumstances during sectioning, there were fewer sections of the VTA with sufficient quality available, which ultimately led to lower numbers of animals in the VTA comparison. The visual representation in Fig. 3 and the similarity of the directionality of the results with those of the NAc, however, do suggest that differences in the VTA may be found in a higher-powered study. Previously, it has been found that a short 3 day high-fat diet does also not change c-Fos expression in the VTA in response to a palatable glucose solution [39], but several reward-related receptors in the VTA are found to be down-regulated after long-term high-calories food diets [51-53]. Unpublished data from our laboratory, gathered with fiber photometry, also suggests that neurons in the VTA become activated upon exposure to a CAF reward in control animals, while these levels of VTA activation are reduced in the context of chronic *ad libitum* CAF diet treatment. The finding that this reduction in VTA activity is more pronounced when exposed to CAF diet then high fat/high sugar pellets, suggests that the element of choice in the CAF diet regime might play a more important role than the nutritional values of the food (unpublished data). Future research is needed to reveal whether different regimes of CAF diet exposure can cause desensitization of VTA neurons.

In addition, we found that different regimes of CAF diet exposure did not change the neural activation of the PFC in response to a food reward. Since the c-Fos-IR cell levels of CTR-CAF rats were also not significantly different from CTR-chow rats in the PFC, we conclude that the PrL and IL of the PFC are not involved in this aspect of reward processing. This is in line with another study that also failed to detect changes in monoamine neurotransmitters in the PFC after CAF exposure [54]. However, this does not mean that the PFC is not affected by junk food at all. It has been shown before, for example, that inbred obesity-prone rats have lower dopamine turn-over in the PFC in response to acute feeding [45]. It is then logic to assume that changes in neural activation within the PFC could be found, if different reward-related aspects were explored. In addition, a small positive correlation of protein intake during the reward exposure resulted in more c-Fos expressing cells in the PrL. This could indicate that proteins could acutely affect the PrL.

At last, we would like to stress that our study was performed in female Long-Evans rats. To the best of our knowledge, this is the first study exploring the effects of chronic junk food exposure on the brain reward system in females. It is important to consider that sex differences can occur and that male and female rats can respond differently to CAF diet. As such, sex differences in changes in reward relevant parameters such the TH mRNA levels and I-opioid receptors expression have already been reported [55]. This highlights the importance of investigating sex differences in the neurobiological response to palatable food intake and the need for more studies in this area being conducted with female rats.

## 5. Conclusion

In conclusion, our study revealed that 6 weeks exposure to CAF diet can decrease c-Fos expression in the NAc-shell when presented with a CAF reward. This decrease was found in all CAF diet regimes: with chronic *ad libitum* and restricted access, and with intermittent (1 day a week) exposure to CAF diet. While no changes in c-Fos expression upon the different diet regimes were found in the PFC, future research should reveal potential decreases in neural activity in the VTA. Our data suggests that long-term junk food exposure can affect the brain reward system, resulting in an attenuated activity of the NAc-shell.

## Acknowledgements

Financial support was received from Helse Nord (HNF1443-19). We thank Carles Baldellou, Laia Oliva and the technical staff of the Unitat d’Experimentació Animal in the Campus Mundet of Universitat de Barcelona (UB) for the excellent care of the animals. Finally, we thank the Advanced Microscopy Core Facility of UiT The Arctic University of Tromsø for access to their equipment for the brain imaging analyses.

## Author contributions

RH: Experimental design, methodology, data gathering, data curation, analysis, supervision, writing – original draft, funding acquisition

JFL: Experimental design, data gathering, writing – review and editing

MS: Methodology, writing - review and editing

AMA: Data gathering

SGA: Data gathering

RME: Methodology, writing - review and editing

EMSS: Experimental design, methodology, data gathering, data curation, analysis, supervision, writing – original draft, funding acquisition

## Notes

### Competing Interest Statement

The authors have declared no competing interest.

### Summary of Updates

Figures were changed, and data on the effects of the nutritional value of the reward on c-Fos expression was added.

