## Supplemental data for "Changes in reward-induced neural activity upon Cafeteria Diet consumption"

**Table S1**. Nutritional information for the standard (STD) chow and the cafeteria (CAF) diet components

| | | CAF diet | | | | | | | |
|---|---|---|---|---|---|---|---|---|---|
| **Nutritional Values (per 100 g)** | *STD chow[(1)]* | *Pork jowl* | *Biscuit with pâté[(2)]* | *Biscuit with cheese[(3)]* | *Muffins* | *Carrots* | *Milk[(4)]* | *Sugar[(4)]* | *Gelatin[(4)]* |
| **Energy (kcal)** | 290.00 | 670.00 | 315.54 | 345.56 | 428.00 | 39,20 | 63.00 | 400.00 | 357.00 |
| **Proteins (g)** | 14.30 | 10.00 | 8.03 | 8.69 | 4.60 | 1.10 | 3.10 | 0.00 | 89.00 |
| **Carbohydrates (g)** | 48.00 | 0.00 | 42.11 | 38.31 | 48.00 | 7.80 | 4.60 | 100.00 | 0.00 |
| **Sugars (g)** | 0.00 | 0.00 | 11.22 | 12.56 | 32.00 | 6.80 | 4.60 | 100.00 | 0.00 |
| **Lipids (g)** | 4.00 | 70.00 | 12.46 | 16.85 | 24.00 | 0.40 | 3.60 | 0.00 | 0.00 |
| **Fiber (g)** | 22.10 | 0.00 | 1.53 | 1.47 | 0.50 | 2.60 | 0.00 | 0.00 | 0.00 |
| **Salt (Na) (g)[5]** | 0.10 | 0.00 | 1.29 | 1.40 | 0.55 | 0.07 | 0.13 | 0.00 | 1,2 |
| **Water (g)** | 0.00 | 18.00 | 26.24 | 30.41 | 28.00 | 87.80 | 88.40 | 0.50 | 12 |

[1] Teklad Global 14% Protein Rodent Diet by Harlan. [2] Nutritional information from digestive biscuits plus liver pork pâté. [3] Nutritional information from digestive biscuits plus semi-spreadable cheese. [4] These products are the ingredients of flam at a proportion of 220 g sugar:1L milk: 20 g gelatin. [5] Values of salt content are presented as sodium (g).
The CAF diet contained the following products: Pork jowl: 50% Tocino de cerdo (fat) and 50% Zanahorias Frascas (belly); Biscuits: Galletas Maria dorada Mercadona; Pâté: Paté de cerdo sabopr suave Argal; Cheese: Week 1-2: Queso en porciones Week 3-6: Caserio Quesilete; Muffins: Magdalenas Hacendado Paquete; Carrots: Zanahorias Frescas; Milk: Leche entera Hacendado; Sugar: Azucar Blanco; Gelatin: Gelatina Neutral.

**Table S2**. Reward food intake (g) during the CAF reward exposure per animal

| Foodstuff / Group | CTR-Chow | CTR-CAF | CAFchr | CAFint | CAFres |
|---|---|---|---|---|---|
| **Chow** | 1.05 ± 0.321 | - | - | - | - |
| **Muffin** | - | 0.03 ± 0.10$^{a}$ | 0.00 ± 0.00$^{b}$ | 0.19 ± 0.06$^{a,b}$ | 0.05 ± 0.02$^{b}$ |
| **Pork jowl** | - | 0.74 ± 0.22 | 0.29 ± 0.07 | 0.65 ± 0.27 | 1.61 ± 0.59 |
| **Biscuit & Pâté** | - | 0.35 ± 0.06$^{a}$ | 0.50 ± 0.06$^{a,b}$ | 0.89 ± 0.20$^{b}$ | 0.38 ± 0.13$^{a}$ |
| **"Flam"** | - | 1.93 ± 0.38$^{a}$ | 1.98 ± 0.38$^{a}$ | 3.99 ± 0.66$^{b}$ | 3.01 ± 0.52$^{a,b}$ |

The data represents the reward intake on the perfusion day when the rats received standard chow or CAF reward
$^{abc}$ Mean ± SEM values within a row with unlike letters were significantly different between groups (one-way ANOVA and Bonferroni's *post hoc* comparison) $p<0.05$, g=grams, "flam" =gelatinized whole milk with sugar. n=8 animals per group.

**Table S3**. Reward food intake of energy and nutrients during the CAF reward exposure per animal

| Nutritional Values / Group | CTR-Chow | CTR-CAF | CAFchr | CAFint | CAFres |
|---|---|---|---|---|---|
| **Energy (kcal)** | 3.05 ± 0.93$^{a}$ | 9.85 ± 1.49$^{a,b}$ | 5.96 ± 0.66$^{a,b}$ | 13.04 ± 2.11$^{a,b}$ | 16.04 ± 4.58$^{b}$ |
| **Proteins (g)** | 0.15 ± 0.02 | 0.17 ± 0.02 | 0.12 ± 0.01 | 0.26 ± 0.04 | 0.28 ± 0.07 |
| **Carbohydrates (g)** | 0.50 ± 0.08$^{a}$ | 0.63 ± 0.07$^{a}$ | 0.67 ± 0.07$^{a}$ | 1.30 ± 0.22$^{b}$ | 0.61 ± 0.09$^{a,b}$ |
| **Sugars (g)** | 0.00 ± 0.00$^{a}$ | 0.56 ± 0.08$^{b}$ | 0.47 ± 0.08$^{b}$ | 1.01 ± 0.14$^{c}$ | 0.70 ± 0.11$^{b,c}$ |
| **Lipids (g)** | 0.04 ± 0.01$^{a}$ | 0.86 ± 0.27$^{a,b}$ | 0.43 ± 0.05$^{a}$ | 0.91 ± 0.36$^{a,b}$ | 0.32 ± 0.12$^{b}$ |
| **Salt (Na) (g)[1]** | 0.001 ± 0.010$^{a}$ | 0.006 ± 0.001$^{a,b}$ | 0.004 ± 0.000$^{a,b}$ | 0.007 ± 0.001$^{b}$ | 0.004 ± 0.001$^{a,b}$ |
| **Water from food (g)** | 0.00 ± 0.00$^{a}$ | 1.99 ± 0.28$^{a}$ | 1.58 ± 0.27$^{a}$ | 12.69 ± 1.92$^{b}$ | 2.59 ± 0.46$^{a}$ |

The data represents the reward intake on the perfusion day when the rats received standard chow or CAF reward.
$^{abc}$ Mean (± SEM) values within a row with unlike letters were significantly different between groups (one-way ANOVA and Bonferroni's *post hoc* comparison) $p<0.05$. kcal=kilocalories, g=grams. n=8 animals per group. [1] Values of salt content are presented as sodium (g).

### Figure S1.

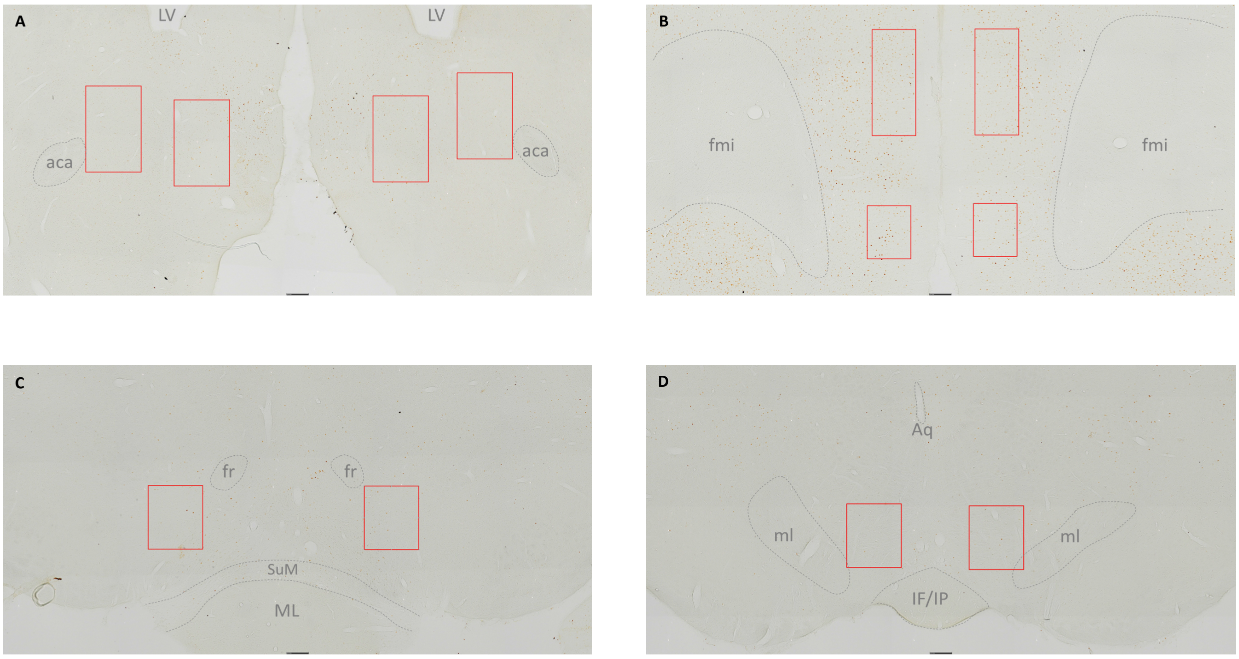


***Figure S1.*** *Representation of the box placement in A) the nucleus accumbens (NAc), B) the prefrontal cortex (PFC), C) the anterior ventral tegmental area (VTA) and D), the posterior VTA with the boxes in red. aca= anterior commissure, Aq= aqueduct, fmi = corpus callosum, forceps minor, fr = fasciculus retroflexus, IF/IP = interfascicular/interpeduncular nucleus, LV = lateral ventricle, ML = mammillary nucleus, Ml = medial lemniscus, SuM = supramammillary nucleus.*
